# ModCRElib: A standalone package to model *cis*-regulatory elements

**DOI:** 10.64898/2026.02.18.701901

**Authors:** Patrick Gohl, Oriol Fornes, Patricia M. Bota, Alberto Meseguer, Jaume Bonet, Ruben Molina-Fernández, Joan Planas-Iglesias, Altair C. Hernandez, Oriol Gallego, Narcis Fernandez-Fuentes, Baldo Oliva

## Abstract

**Summary:** The ModCRElib package provides various tools for the analysis and modelling of transcription factor(TF)-DNA and regulatory complex inter-protein interactions. It takes structural information on these interactions to predict TF binding motifs, generate binding profiles along DNA sequences that score the binding affinity, predict TF binding sites and model the structure of higher order regulatory complexes. It is capable of working with a variety of input data formats and sources. The user may follow the analysis pipeline as outlined in the documentation or the user can make use of any of the multiple functionalities in an isolated manner. The package takes the service offered by the ModCRE server and enables users to apply its tools in an unrestricted and customizable manner. In this paper we provide 5 example uses of ModCRElib. This includes (i) TF binding affinity prediction, (ii) TF binding aggregation, (iii) characterization of specificity in TF binding sites along target DNA sequences,(iv) modelling TF bound to predicted binding sites, and (v) the generation of statistical potential derived scoring profiles of TF interacting with DNA.

**Availability:** https://github.com/structuralbioinformatics/ModCRElib

**Supplementary information:** Available at https://github.com/structuralbioinformatics/ModCRElib. doi:10.5281/zenodo.17484081

## 1 Introduction

With the advent of deep learning based protein fold predictors such as AlphaFold3 (Abramson et al. 2024) and RoseTTAFold (Baek et al. 2021) the field of reliable protein structural prediction has expanded from the scope of template based methods such as Modeller (Webb and Sali 2016) to include proteins and protein complexes that could not be predicted by comparative modeling alone.. This growing database of protein structural information is a unique source of knowledge requiring only the development of tools to exploit this knowledge for real world applications. One such possible application is the analysis of transcriptional regulatory elements. As a key part of the regulation of gene expression, transcriptional regulatory elements contain transcription factor (TF) binding sites which, upon TF-binding, can up or downregulate the synthesis of associated gene products (Andersson & Sandelin 2020). These gene products, or lack thereof, impact cell development and homeostasis as well as the response to diseases and many more biological functions reliant on proteins or RNA. In this context, recent advances in AI have introduced applications capable of characterizing the specificity of transcription factor binding solely from their structural information (Mitra et al. 2024).

Building on the development of methodology to apply statistical potentials to evaluate protein interactions (Feliu et al. 2011, Fornes et al. 2014, Meseguer et al. 2020) we created ModCRE (Fornes et al. 2024), a webserver applying statistical potentials derived from structural models to evaluate and predict the interactions of TF with DNA, co-factors and other TFs. We chose to apply statistical potentials for their ability to incorporate non-structural experimental information as well as the recent development of better tools for the prediction of TF structures. ModCRE is capable of taking user input in the form of amino acid or DNA sequences as well as protein structures to predict TF position weight matrices (PWM) which it uses to predict TF binding sites. It may also predict structural models of TF interactions with co-factors and other TFs as well as generate and compare TF binding profiles along target DNA sequences that score the affinity of the TF per nucleotide or amino-acid position (see an example in de Martin et al 2024). However, its distribution as a webserver, while keeping it as user friendly as possible, limited its use in scope and depth (restrictions on scannable DNA length, limited job queuing etc.). To address these limitations we developed a ModCRE standalone python package: ModCRElib. ModCRElib has been outfitted with all of the tools available on the ModCRE web server plus additional applications (see GitHub repository). Additionally, any necessary database for the running of these tools have been made freely accessible for download. A GitHub repository containing detailed instructions on download and use was prepared with examples outlining steps to take in ModCRElib to replicate the functionality of ModCRE and illustrate additional processing and analysis steps to build on what ModCRE has provided until now.

## 2 ModCRElib Architecture

### Installation

The ModCRElib library can be installed by cloning the GitHub repository. Installation instructions regarding dependencies and specific terminal commands to follow along with are outlined in the repositories README. After installation, “PBM” and “PDB” folders need to be retrieved from http://sbi.upf.edu/modcrefiles/, or generated locally using ModCRElib scripts. After installation we provide the user with 7 bash scripts that can be called to start running either on the input examples provided or directly applied to user data.

### Input

The required input varies and depends on the level of analysis and pipeline entry point required. To begin at the most basic level, the user may simply execute a script to search for TF sequences from the Uniprot (The Uniprot Consortium 2025) or the Protein Data Bank (PDB) (Berman et al. 2000) databases. If, on the other hand, they wish to proceed with specific TFs they would need to provide a single fasta or multi-fasta file containing the full amino acid sequences of those TFs. In either case one or more structures of the provided TF interacting with DNA will be modelled. This lays the foundation for all analysis possible within ModCRElib, with the additional requirement of DNA sequences which must be provided either as a fasta file or string parameter at various stages within the pipeline.

### Integration of external software

The library requires and can handle the results from external software. Most of these dependencies are inherited from ModCRE. A list of software dependencies has been provided on GitHub (in a README file). External software must be installed locally (with the exception of SBILib (Gohl et al. 2023), a version of which is provided with ModCRElib), and users can manually specify their system address in the configuration file (ModCRElib/configure/config.ini).

## 3 ModCRElib capabilities

### TF binding specificity prediction

The prediction of TF-DNA binding specificity within ModCRElib requires one 3D structure of the target TF binding to DNA, preferably multiple predicted structures. These models may be derived from ModCRElib predictions, PDB models, or any other method generating protein structures such as BioEmu (Lewis et al. 2025). The TF-DNA complex is then used to assign scores to all possible DNA sequences of the binding site using statistical potentials. Then an alignment of the top scoring DNA sequences is used to make a PWM prediction as outlined in (Fornes et al. 2024). The output is stored in a user defined folder where the PWM may be viewed in .pwm or .meme format as well as plots of the forward and reverse logo of the pwm.

### PWM Aggregation

ModCRElib is designed to generate multiple PWM predictions for each TF based on structural models of TF–DNA interactions. For a given TF sequence, several structural models can be constructed using different templates from the PDB, resulting in multiple PWM predictions. The degree of similarity among these predictions depends on the variability of the selected templates and on the specific regions of the TF sequence that are modeled. This landscape of PWMs is intended to capture the multi-patterned binding of TF, the finer details of which are lost when they are forced into a single representative PWM. Through its aggregation function ModCRElib allows a user to aggregate predicted PWMs from an input folder into clusters. It will then generate a single representative PWM for each cluster and store these along with cluster member PWMs for the user. Thus, ModCRElib enables users to proceed with downstream regulation analysis with a more fine grain TF binding affinity representation.

### Scanning DNA for binding sites

When scanning a DNA sequence for potential TF-specific recognized sequences used as binding sites, the user may opt to use the provided database of PWMs or provide their own selection to scan with. In either case ModCRElib will use FIMO (Grant et al. 2011) to scan the DNA sequence for hits passing the user defined p-value threshold. ModCRElib will then automatically generate thread files for each hit (i.e. TF-DNA binding site).

### Modelling regulatory complexes

In order to start the modelling process the user will first have to generate a folder containing the coordinates of binary interactions in standard PDB format. These binary interaction files will consist of TF-DNA interactions in the case of a TF binding site or TF-TF/TF-cofactor interactions in the case of protein-protein interactions. The TF-DNA interactions may be retrieved by generating pdb files from the output of ModCRElib’s DNA scanning function. In the case of the protein-protein interactions the files will have to be retrieved from external sources such as CM2D3 (Bota et al. 2023), Modpin (Meseguer et al. 2020), or AlphaFold3. Lastly, a structural model of the DNA sequence will have to be included in the folder. ModCRElib will then use the folder of binary interactions to iteratively attempt superimposing various structures (while avoiding clashes) to generate the largest possible complex.

### Generating scoring profile of TF-DNA binding

Scoring profiles can be generated to analyse the binding of a TF along a DNA sequence, check the effects of Single Nucleotide Polymorphisms (SNPs) on the binding of a TF or compare the binding of different TFs along the same DNA sequence. In order to generate these profiles the user will have to provide ModCRElib with structural models of the TF interacting with a DNA double-helix (which may be created with the modelling function). ModCRElib can generate a profile from a single model, but it is best used in conjunction with multiple predicted structures per tested TF. As with the PWM prediction, the DNA sequence will be fed through the model and have statistical potentials assigned at each position. If the user provided multiple models, the mean of the profile will then be calculated and saved as a csv table. The script can either generate a plot automatically (Fig. 1) or a simple python script available within ModCRElib can be used to plot the profile for visualization purposes.

**Fig. 1.**
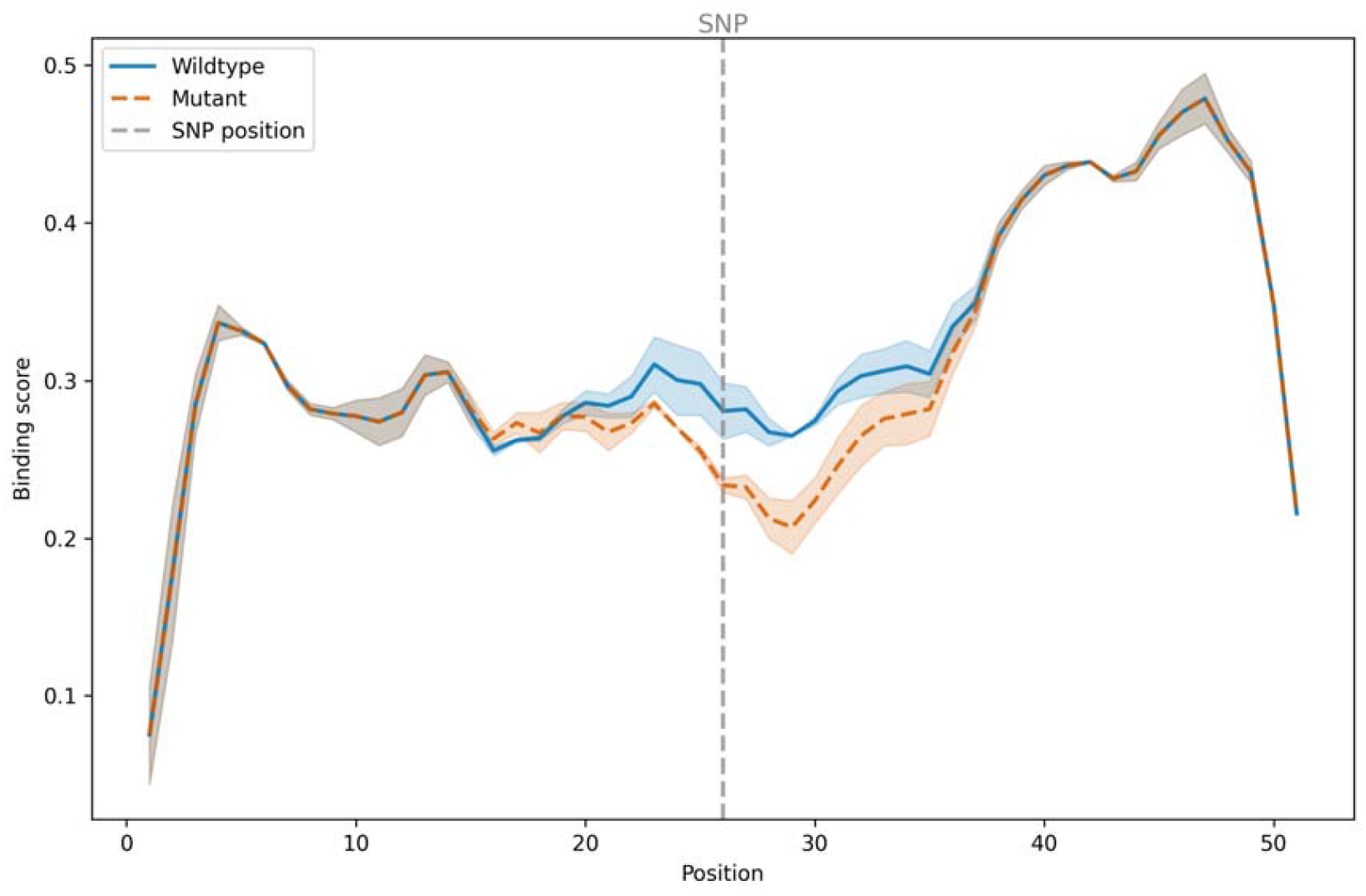
Illustration of TF-DNA energy-binding profile measured by normalized statistical potentials with ModCRElib. An allele specific binding event for the AHR TF was selected (rs573372954). Both the mutant (orange) and wildtype (blue) DNA-binding sequences and two models generated for AHR were submitted for profiling. The mutation is located at DNA position 26. Shaded areas represent standard error. Visible is the drop in binding score for the mutant sequence reflecting the allele specific binding event.

## 1 Examples

A detailed step-by-step guide outlining the execution of both the aforementioned examples as well as other applications has been made available in the GitHub repository. The GitHub repository provides additional support in handling the various parameters for further customization of ModCRElib jobs as well as information on how to run jobs in parallel.

## 5 Conclusion

ModCRElib is an open-source library for the analysis and modelling of *cis*-regulatory complexes. It provides all of the functionality of ModCRE without any of the restrictions levied upon it as a webserver. ModCRElib provides additional functionality not available to ModCRE users by enabling the modulation of jobs, extraction and separate application of constituent scripts as well as the introduction of new scripts with new functionalities. ModCRElib will provide the growing community of ModCRE users with additional functionality and flexibility in meeting their needs.

## Funding

The work was supported by grants PID2020-113203RB-I00 and “Unidad de Excelencia María de Maeztu” (ref: CEX2018-000792-M), funded by the MCIN and the AEI (DOI: 10.13039/501100011033) as well as an FPU scholarship (ref: FPU22/02303) and an SGR from the Generalitat de Catalunya (ref: 4413015318-J.SELENT/SGR-22). OG acknowledges PID2021-127773NB-I00 from AEI. JPI acknowledges the European Union’s Horizon Europe Framework Programme under the grant agreement No. 101136607 (CLARA), and e-INFRA CZ and ELIXIR-CZ projects (LM2023055 and LM2018140), supported by the Ministry of Education, Youth and Sports of the Czech Republic, for the computational resources provided.

The authors thank the Scientific Computing Core Facility (MELIS-UPF).

## References

Abramson, J., Adler, J., Dunger, J., Evans, R., Green, T., Pritzel, A., … & Jumper, J. M. (2024). Accurate structure prediction of biomolecular interactions with AlphaFold 3. Nature, 630(8016), 493–500.

Baek, M., DiMaio, F., Anishchenko, I., Dauparas, J., Ovchinnikov, S., Lee, G. R., … & Baker, D. (2021). Accurate prediction of protein structures and interactions using a three-track neural network. Science, 373(6557), 871–876.

Andersson, R. & Sandelin, A. Determinants of enhancer and promoter activities of regulatory elements. Nat Rev Genet 21, 71–87 (2020)

Berman, H. M., Westbrook, J., Feng, Z., Gilliland, G., Bhat, T. N., Weissig, H., … & Bourne, P. E. (2000). The protein data bank. Nucleic acids research, 28(1), 235–242.

Bota, P. M., Hernandez, A. C., Segura, J., Gallego, O., Oliva, B., & Fernandez-Fuentes, N. (2023). CM2D3: furnishing the human interactome with structural models of protein complexes derived by comparative modeling and docking. Journal of Molecular Biology, 435(14), 168055.

de Martin, X., Oliva, B., & Santpere, G. (2024). Recruitment of homodimeric proneural factors by conserved CAT–CAT E-boxes drives major epigenetic reconfiguration in cortical neurogenesis. Nucleic Acids Research, 52(21), 12895–12917.

Feliu, E., Aloy, P., & Oliva, B. (2011). On the analysis of protein–protein interactions via knowledge□based potentials for the prediction of protein–protein docking. Protein Science, 20(3), 529–541.

Fornes, O., Garcia-Garcia, J., Bonet, J., & Oliva, B. (2014). On the use of knowledge-based potentials for the evaluation of models of protein–protein, protein–DNA, and protein–RNA interactions. Advances in protein chemistry and structural biology, 94, 77–120.

Fornes, O., Meseguer, A., Aguirre-Plans, J., Gohl, P., Bota, P. M., Molina-Fernández, R., … & Oliva, B. (2024). Structure-based learning to predict and model protein–DNA interactions and transcription-factor co-operativity in cis-regulatory elements. NAR Genomics and Bioinformatics, 6(2), lqae068.

Gohl, P., Bonet, J., Fornes, O., Planas-Iglesias, J., Fernandez-Fuentes, N., & Oliva, B. (2023). SBILib: a handle for protein modeling and engineering. Bioinformatics, 39(10), btad613.

Grant, C. E., Bailey, T. L., & Noble, W. S. (2011). FIMO: scanning for occurrences of a given motif. Bioinformatics (Oxford, England), 27(7), 1017–1018. 10.1093/bioinformatics/btr064

Lewis, S., Hempel, T., Jiménez-Luna, J., Gastegger, M., Xie, Y., Foong, A. Y., … & Noé, F. (2025). Scalable emulation of protein equilibrium ensembles with generative deep learning. Science, 389(6761), eadv9817.

Meseguer, A., Årman, F., Fornes, O., Molina-Fernández, R., Bonet, J., Fernandez-Fuentes, N., & Oliva, B. (2020). On the prediction of DNA-binding preferences of C2H2-ZF domains using structural models: application on human CTCF. NAR Genomics and Bioinformatics, 2(3), lqaa046.

Meseguer, A., Dominguez, L., Bota, P. M., Aguirre□Plans, J., Bonet, J., Fernandez□Fuentes, N., & Oliva, B. (2020). Using collections of structural models to predict changes of binding affinity caused by mutations in protein–protein interactions. Protein Science, 29(10), 2112–2130.

Mitra, R., Li, J., Sagendorf, J. M., Jiang, Y., Cohen, A. S., Chiu, T. P., … & Rohs, R. (2024). Geometric deep learning of protein–DNA binding specificity. Nature Methods, 21(9), 1674–1683.

UniProt: the universal protein knowledgebase in 2025. Nucleic acids research, 2025, 53.D1: D609–D617.

Webb, B., & Sali, A. (2016). Comparative protein structure modeling using MODELLER. Current protocols in bioinformatics, 54(1), 5–6.

